# Integrating Transcriptomics, Genomics, and Imaging in Alzheimer’s Disease: A Federated Model

**DOI:** 10.1101/2021.09.14.460367

**Authors:** Jianfeng Wu, Yanxi Chen, Panwen Wang, Richard J Caselli, Paul M Thompson, Junwen Wang, Yalin Wang, for the Alzheimer’s Disease Neuroimaging Initiative

**Author notes:** **Please address correspondence to:** Dr. Junwen Wang, Department of Health Sciences Research & Center for Individualized Medicine, Mayo Clinical Arizona, 13400 E Shea Blvd, Scottsdale, AZ 85259 USA, **E-mail:**, Dr. Yalin Wang, School of Computing and Augmented Intelligence, Arizona State University, P.O. Box 878809, Tempe, AZ 85287 USA, **E-mail:**. **Authors contributed equally** (Junwen Wang, Yalin Wang). Acknowledgments: Data used in preparing this article were obtained from the Alzheimer’s Disease Neuroimaging Initiative (ADNI) database (adni.loni.usc.edu). As such, many investigators within the ADNI contributed to the design and implementation of ADNI and/or provided data but did not participate in the analysis or writing of this report. A complete listing of ADNI investigators can be found at: http://adni.loni.usc.edu/wp-content/uploads/how_to_apply/ADNI_Acknowledgement_List.pdf.

## Abstract

Alzheimer’s disease (AD) affects more than 1 in 9 people age 65 and older and becomes an urgent public health concern as the global population ages. In clinical practice, structural magnetic resonance imaging (sMRI) is the most accessible and widely used diagnostic imaging modality. Additionally, genome-wide association studies (GWAS) and transcriptomics – the study of gene expression – also play an important role in understanding AD etiology and progression. Sophisticated imaging genetics systems have been developed to discover genetic factors that consistently affect brain function and structure. However, most studies to date focused on the relationships between brain sMRI and GWAS or brain sMRI and transcriptomics. To our knowledge, few methods have been developed to discover and infer multimodal relationships among sMRI, GWAS, and transcriptomics. To address this, we propose a novel federated model, Genotype-Expression-Imaging Data Integration (GEIDI), to identify genetic and transcriptomic influences on brain sMRI measures. The relationships between brain imaging measures and gene expression are allowed to depend on a person’s genotype at the single-nucleotide polymorphism (SNP) level, making the inferences adaptive and personalized. We performed extensive experiments on publicly available Alzheimer’s Disease Neuroimaging Initiative (ADNI) dataset. Experimental results demonstrated our proposed method outperformed state-of-the-art expression quantitative trait loci (eQTL) methods for detecting genetic and transcriptomic factors related to AD and has stable performance when data are integrated from multiple sites. Our GEIDI approach may offer novel insights into the relationship among image biomarkers, genotypes, and gene expression and help discover novel genetic targets for potential AD drug treatments.

## 1 INTRODUCTION

Alzheimer’s disease (AD) is a major public health concern, with the number of affected individuals expected to triple, reaching 13.8 million, by the year 2050 in the U.S. alone (Brookmeyer et al., 2007). Current therapeutic failures in patients with dementia due to AD may be due to interventions that are too late or targets that are secondary effects and less relevant to disease initiation and early progression (Hyman, 2011). Mounting evidence suggests that germline mutations, e.g., DNA single nucleotide polymorphisms (SNPs), play an important role in AD etiology and progression (Kunkle et al., 2019; Murrell et al., 1991). Among various genetic risk factors, Apolipoprotein E (*APOE*) has the strongest association to late-onset AD, and the e4 allele is associated with increased risk, whereas the e2 allele is associated with decreased risk (Bertram et al., 2007). Known genetic risk variants could be used to identify presymptomatic individuals at risk for AD and support diagnostic assessment of symptomatic subjects. By taking into account patients’ genetic risk factors, at-risk individuals could be more readily identified, diagnostic precision could be improved, and targetable disease mechanisms for new drug development may be discovered (Freudenberg-Hua et al., 2018; Lambert et al., 2013; Mormino et al., 2016; Singanamalli et al., 2017). By enabling each patient to receive earlier diagnoses, risk assessments, and optimal treatments, personalized or precision medicine holds promise for improving early AD intervention while also lowering costs (Vogenberg et al., 2010).

Recent clinical trials targeting single molecular mechanisms have failed (Cummings et al., 2014; Mehta et al., 2017). Rather, it might be necessary to tackle the problem from a holistic or multimodality perspective (Pimplikar, 2017; Xicota et al., 2019). Indeed, the NIH and the scientific community realized this problem a while ago and have already started to produce multi-omics data. For example, the Alzheimer’s Disease Sequencing Project (ADSP) data repository contains genomic level data derived from genome-wide association studies (GWAS) (Kunkle et al., 2019; Saykin et al., 2015), whole-exome sequencing (WES) (Bis et al., 2020; Simino et al., 2017), and whole-genome sequencing (WGS), and RNA level data including mRNA, miRNA, and long non-coding RNA profiling from either microarray or RNA-Seq (Piras et al., 2017). And transcriptome-wide association studies (TWASs) provides a way to use eQTLs and expression data to guide GWAS of AD (Luningham et al., 2020). Brain imaging has played a significant role in the study of Alzheimer’s disease (Johnson et al., 2012). Integrating imaging data and omics data is becoming an emerging data science field known as imaging genomics (Shen and Thompson, 2020). The major task of this field is to perform integrated analysis of imaging and omics data, often combined with other biomarkers, as well as clinical and environmental data. The ultimate goal is to gain new insights into the underlying mechanisms of human health and disease, to better inform the development of new diagnostic, therapeutic, and preventative approaches.

Various imaging genetics methods have been developed to integrate imaging and genetic data. However, most studies have focused on imaging, imaging combined with GWAS data (Chauhan et al., 2015; Grasby et al., 2020; Li et al., 2017), imaging with transcriptomics (Ritchie et al., 2018), or GWAS with transcriptomics (Albert and Kruglyak, 2015). For example, imaging genetics has been used to link SNPs with image features (Stein et al., 2010), and expression quantitative trait loci (eQTL) have been used to discover *APOE*-related genes (Zhang et al., 2018). However, relatively few methods (Lv et al., 2017) have been developed to integrate GWAS/WES/WGS, imaging, and transcriptomic data to infer multimodal relationships. Such a multimodal approach may give us a more holistic view of the evidence from multiple sources to provide novel insights on the molecular mechanisms of AD pathogenesis and prognosis. Besides, both gene expression and imaging features are dynamic and change with time and throughout the disease, whereas germline SNPs are unchanged over an individual’s lifetime. We need a better model for studying SNP-image-gene expression relationships to consider both the dynamic changes in imaging and gene expression features and understand how they are affected by an individual’s SNPs. Such knowledge will provide novel insights into the relationship among image biomarkers, genotypes and gene expression, and may help to discover novel genetic targets for pharmaceutical interventions.

AD is a complex multifactorial disorder that involves many biological processes. The launch of the Alzheimer Precision Medicine Initiative (APMI) and its associated cohort program in 2016— facilitated by the academic core coordinating center run by the Sorbonne University Clinical Research Group in Alzheimer Precision Medicine—is intended to improve clinical diagnostics and drug development research in Alzheimer’s disease (Hampel et al., 2019). Hampel et al. (2019) indicate the challenges for precision medicine, including secure data access accompanied by rigorous privacy protection and the availability of data to qualified researchers who may use them to exercise their creative thinking with an *a posteriori* approach or to test their *a priori* hypotheses. Integrating data from multiple sites and sources is common practice to achieve larger sample sizes and to increase statistical power. Unprecedentedly large amounts of biomedical data now exist across hospitals and research institutions. However, different institutions may not be readily able to share biomedical research data due to patient privacy concerns, data restrictions based on patient consent or institutional review board (IRB) regulations, and legal complexities; this can present a major obstacle to pooling large scale datasets to discover and understand AD-related genetic information. To remedy this distributed problem, a large-scale collaborative network, ENIGMA consortium, was built (Thompson et al., 2020). Federated learning is an important direction of interest in multi-site neuroimaging research; the use of distributed computing offers an approach to learn from data spread across multiple sites without having to share the raw data directly or to centralize it in any one location. Even so, most ENIGMA and other GWAS studies currently focus on the influence of genetic variants on human brain structures (Chauhan et al., 2015; Hibar et al., 2015; Satizabal et al., 2019; Zhao et al., 2021) or functional measures (Smit et al., 2018) and relatively few have studied the relationships among image biomarkers, genotypes, and gene expression. In this work (Liu et al., 2017), the authors use a brain-wide gene expression profile available in the Allen Human Brain Atlas (AHBA) as a 2-D prior to guide the brain imaging genetics association analysis. Their transcriptome-guided SCCA (TG-SCCA) framework incorporates the gene expression information into the traditional SCCA model.

In this paper, we propose a novel Federated Genotype-Expression-Image Data Integration model (GEIDI) based on the Chow test (Chow, 1960). The intuition behind our multi-omics framework is illustrated in **Figure 1**. Some important image-expression relationships (correlations) may be diluted when the population is mixed together. Still, when we stratify the population based on their genotypes (a gene like *APOE* or a SNP like *rs942439*), we can observe strong correlations (AA and BB groups) across subgroups. Accordingly, as shown in **Figure 2**, our model is designed to detect if the relationships between X (imaging biomarker) and Y (gene expression) are different among the subgroups. The *p*-value of the model is then used to prioritize the trios (genotype-expression-image).

**Figure 1.**
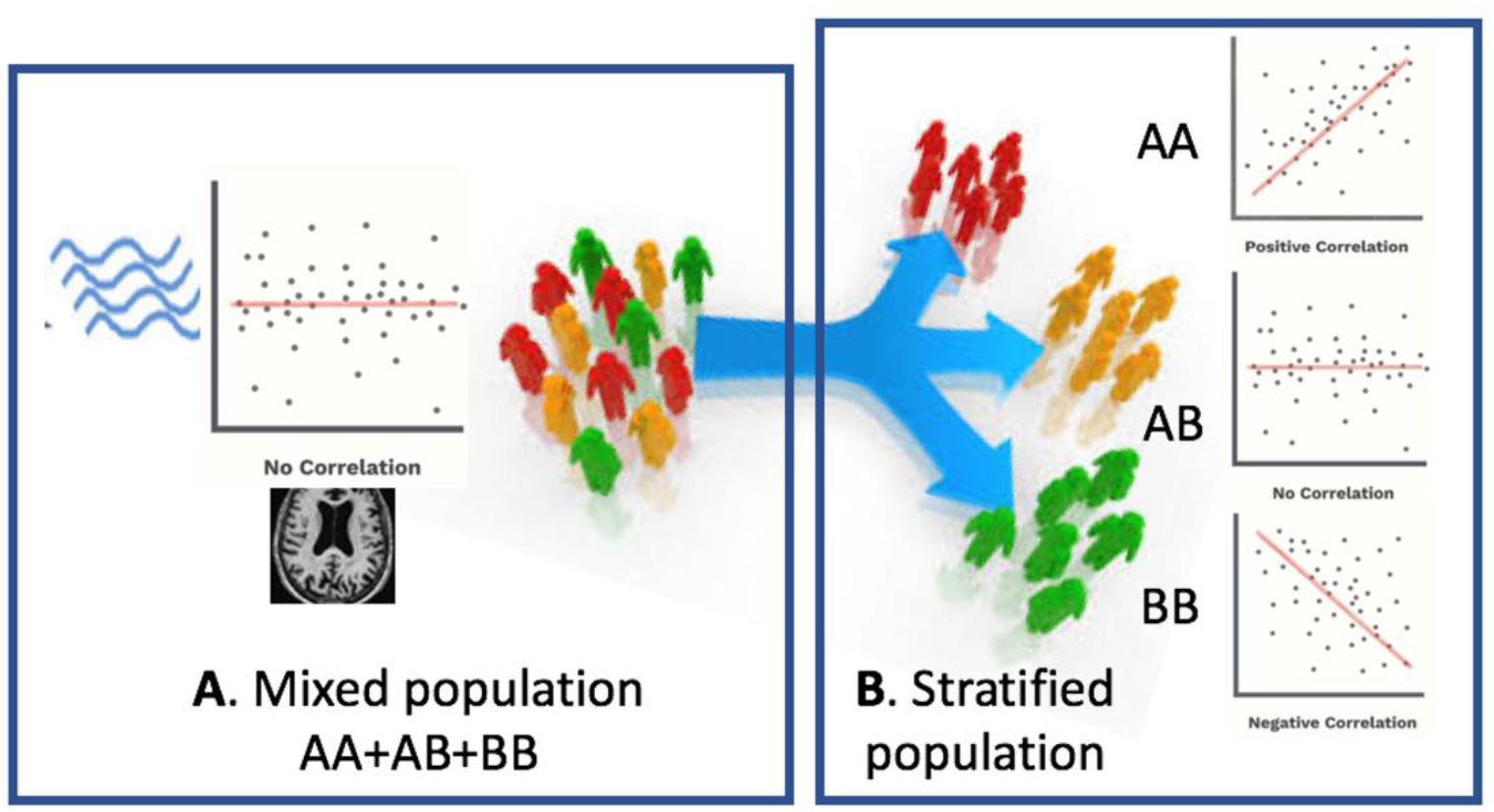
Schematic view of our multi-omics approach. A) When patients are mixed together, image-expression correlation may be low. B) When a certain genotype stratifies patients, some subsets (AA, BB) have a high correlation

**Figure 2.**
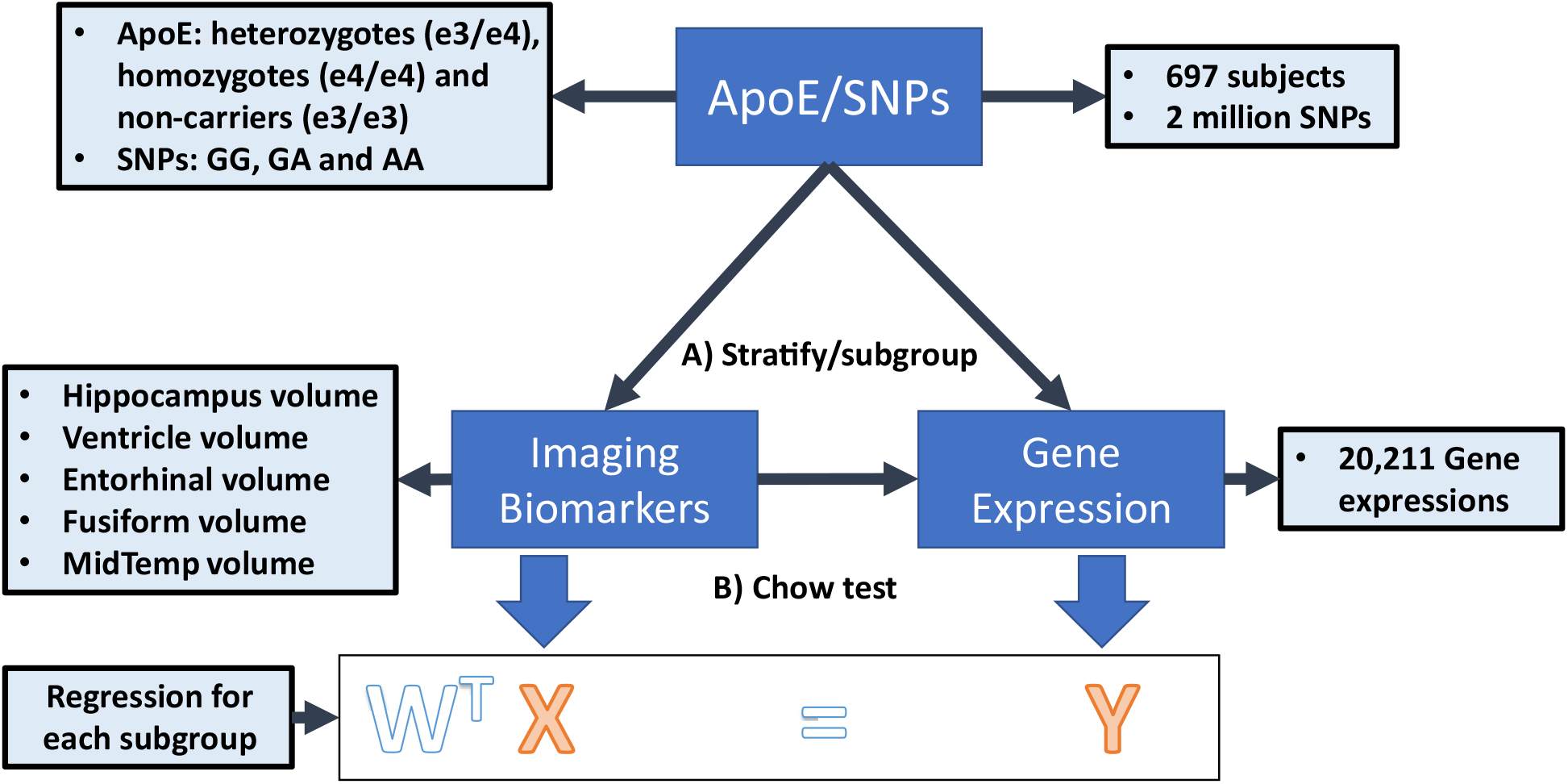
The federated GEIDI model on ADNI data. **A)** Stratify samples into subgroups with different genotypes of a gene (e.g., *APOE*) or at a specific SNP locus (e.g., *rs942439*) **B)** Federated GEIDI is used to detect if the relationships between X (imaging biomarker) and Y (gene expression) are different among the subgroups. The *p*-value of federated GEIDI will then be used to prioritize the trios (genotype-expression-image).

We further design various experiments on publicly available data from the Alzheimer’s Disease Neuroimaging Initiative (ADNI, adni.loni.usc.edu) to demonstrate that our model may detect the genetic factors most related to AD better than the state-of-the-art Matrix eQTL. The overall intent of the work is to detect relationships that inform the design or repurposing of drugs to target these subgroups to achieve precision medicine. We first use a hypergeometric analysis and an AD-related gene list from alzgene.org to evaluate the ability of our federated GEIDI model to discover AD-related gene expression. To further aid in the discovery of genes that may be potential AD drug targets, we also use Pearson correlations analyses to demonstrate the divergence in stratified populations. Additionally, we design experiments to show that our model can discover more AD-related SNPs, based on tests with 1,217 known AD-associated SNPs and 1,217 randomly selected SNPs. Finally, we evaluate the stability of our model under different multi-site conditions. With the ADNI dataset, we set off to test our hypothesis that the proposed federated GEIDI model may be an effective federated model that can provide novel insights into the relationship among image biomarkers, genotypes, and gene expressions and the discovery of novel genes for potential AD drug targets.

## 2 DATA and METHODS

### 2.1 Data preprocessing

The data in this work are from the Alzheimer’s Disease Neuroimaging Initiative (ADNI, adni.loni.usc.edu) and the TADPOLE challenge (tadpole.grand-challenge.org) (Marinescu et al., 2020). The ADNI was launched in 2003 as a public-private partnership led by Principal Investigator Michael W. Weiner, MD. The primary goal of ADNI has been to test whether serial MRI, PET, other biological markers, and clinical and neuropsychological assessments can be combined to measure the progression of MCI and early AD. The genome-wide association study of ADNI is designed to provide researchers with the opportunity to combine genetics with imaging and clinical data to help investigate the mechanisms of the disease. For up-to-date information, see adni.loni.usc.edu/data-samples/data-types/genetic-data/. From the ADNI GWAS, we analyzed data from 697 subjects, including AD patients, people with mild cognitive impairment (MCI), and cognitively unimpaired (CU) subjects, for whom the demographic information is shown in **Table 1**. Each sample has three types of modalities of data: genotypes of known AD risk genes (e.g., *APOE*) and SNPs from genome-wide association studies (GWAS), gene expression measurements (for 20,211 genes) from microarray-based transcriptomic profiling of samples’ blood, and imaging biomarkers from structural magnetic resonance imaging (sMRI) data of subjects’ brains. We use *plink* to perform a quality check of the genotype data. The SNPs in the normal group that deviate significantly from Hardy-Weinberg equilibrium are removed (Purcell et al., 2007). The LINNORM package (Yip et al., 2017) was adopted to perform data transformation on the expression data for normality and homoscedasticity. Recent evaluations (Huang et al., 2018; Yip et al., 2018) show that LINNORM typically performs better than current DEG analysis methods for both single-cell and bulk RNA-Seq, such as Seurat (Satija et al., 2015) and DESeq2 (Love et al., 2014).

**Table 1.** Demographic information for the subjects we study from the ADNI.

| Group | Sex (M/F) | Age | MMSE |
| --- | --- | --- | --- |
| AD (n=96) | 59/37 | 74.8 ± 7.5 | 21.8 ± 4.1 |
| MCI (n=366) | 209/157 | 72.0 ± 7.5 | 28.0 ± 1.7 |
| CU (n=235) | 115/120 | 74.4 ± 5.8 | 29.1 ± 1.2 |

Values are mean ± standard deviation, where applicable.

Eventually, we get 2,059,586 SNPs, *APOE* genotype, and expression data for 20,211 genes for each sample. Besides, from the TADPOLE challenge, we obtained five brain imaging biomarkers for each subject calculated using FreeSurfer (Fischl et al., 1999) with sMRI, including the volume of the hippocampus and middle temporal gyrus (MidTemp). To adjust for individual differences in head size, the volume of each sub-cortical region is adjusted by the intracranial vault volume (ICV) of each subject (volume/ICV). The difference between the date for gene expression collection and MRI scan is less than five months.

### 2.2 Federated genotype-expression-image data integration framework

Econometrician Gregory Chow first proposed the Chow test in 1960 (Chow, 1960) to determine whether correlation coefficients estimated in two subgroups are significantly different. In econometrics, it is most commonly used in time series analysis to test for the presence of a structural break at a period that can be assumed to be known as *a priori* (for instance, a significant historical event such as a war). For example, we can model the data as *y* = *wX* + *ϵ*. Then, the data can be broken into two groups according to some event and fitted to the regression model as, *y*_1_ = *w*_1_*x*_1_ + *ϵ* and *y*_2_ = *w*_2_*x*_2_ + *ϵ*. The null hypothesis of the Chow test asserts that *w*_1_ = *w*_2_ and the model errors *ϵ* are independent and identically distributed from a normal distribution with unknown variance. Let *S*_*C*_, *S*_1_, and *S*_2_ be the sum of squared residuals for the three regression models respectively, *N*_1_ and *N*_2_ are the number of observations in each group, and *k* is the number of parameters. The Chow test statistic is 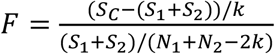, which follows the *F*-distribution with *k* and *N* + *N* ‒ 2*k* degrees of freedom.

Although the Chow test is commonly used in the financial industry, it has not been used in the biomedical field. In this work, we first generalize the Chow test model to estimate the multi-subgroup condition and further introduce a federated learning technique to the model. We apply the proposed model to the ADNI dataset to detect the significant trios among genotype, gene expression, and imaging biomarkers and discover the dominant genetic and transcriptomics factors for brain structures.

#### 2.2.1 Standardization

We simulate the multi-site condition by separating all the samples into *I* hypothetical institutions (*I* = 5) on Apache Spark (spark.apache.org), a state-of-the-art distributed computing platform. (Although the ADNI data can be centralized, such a federated analysis would allow the method to be scaled up to much larger datasets, including genomic data that is difficult to centralize for logistic or regulatory reasons). As illustrated in **Figure 2**, the samples in each institution can be further partitioned into at most three subgroups (*g* = 1,2,3) according to the subject’s genotype at certain SNP loci (e.g. GG, GA, AA) or a gene (e.g., stratified by the three *APOE* genotypes considered in this study). Accordingly, 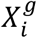 and 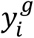 respectively represent the image biomarkers and gene expression values in the *g*th group of the *i*th institution. The data from the *g*th group in all *I* institutions will be fitted into a regression model in a federated strategy.

#### 2.2.2 Federated Chow test analysis

Using federated linear regression, we can calculate four linear models for all the *I* institutions, including three models for three subgroups and one for all samples in the three subgroups. 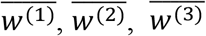.and 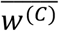 are their optimal coefficient vectors. The Chow test assumes that the errors ϵ Are independent and identically distributed from a normal distribution by an unknown variance. The null hypothesis of the Chow test asserts that 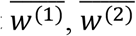, and 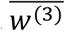 are equal. The predictive test suggested by Chow is then:

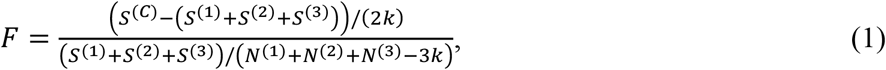

where *S*^(*C*)^ is the sum of squared residuals from the combined data from the three subgroups, *S*^(1)^ is the sum of squared residuals from the first group, and so on for *S*^(2)^ and *S*^(3)^. *N*^(1)^, *N*^(2)^, and *N*^(3)^ are the number of samples in each subgroup, and *k* is the number of parameters. Under the null hypothesis, the test statistic follows the *F*-distribution with 2*k* and *N*^(1)^ + *N*^(2)^ + *N*^(3)^ ‒ 3*k* degrees of freedom. The global center will calculate *F* by gathering all the least square losses and the number of subjects for each subgroup and combined data from each institution. For example, for the first subgroup, the global least-square loss is 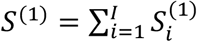 and the global subject number is 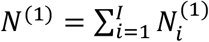. Eventually, the *p*-value will be calculated at the global coordinating center and assigned to each institution.

#### 2.2.3 Federated linear regression

Many regression models may be selected for the Chow test model, such as linear regression (Barbur et al., 1994), polynomial regression (Rawlings et al., 1998), ridge regression (Hoerl and Kennard, 1970), and so on. In this study, we focus on studying the differences in the relationships between imaging biomarkers and gene expression among different groups. Complex regression models, like polynomial regression, may lead to over-fitting and meaningless results. Also, sparse or penalized regression methods, such as ridge regression, require an appropriate regularization parameter. Therefore, in this work, linear regression would be the most rational choice.

Since the federated regression models for each subgroup are the same, we omit the group superscripts here. For the data in one subgroup of all the *I* institutions, we can calculate the linear regression equation as: *y* = *Xw* + ϵ, where *X* ∈ *R*^*N*×*k*^ represents the independent variables, *y* ∈ *R*^*N*^ is a vector of the observations on a dependent variable, *w* ∈ *R*^*k*^ is a coefficient vector, and ϵ ∈ *R*^*N*^ is the disturbance vector. *N* is the number of observations in the group, and *k* is the number of parameters. Then, the coefficient vector *w* can be estimated by minimizing the least squared function, 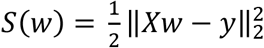.

To avoid centralizing the data, (*X*_*i*_, *y*_*i*_), from each institution, we first rewrite the minimization problem as, 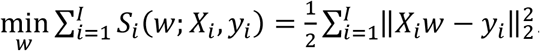. Then, the global gradient can be calculated as, 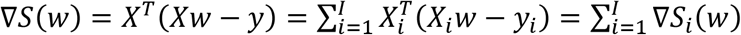. Therefore, instead of centralizing the data, the global center only needs to gather the partial gradient, ∇*S*_*i*_(*w*), which is calculated with (X_i_, y_i_) at each local institution. After computing the global gradient, ∇*S*(*w*), the global center will send it back to *i*th local institution. Finally, *w* will be updated at each institution by gradient descent with the same learning rate, *w* ← *w* ‒ η∇*S*(*w*). The reason for not updating *w* at the global center is to avoid possible data reconstruction. When *w* is zero, the local gradient sent to the center is 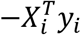. Then, the global center can easily acquire 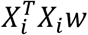 and *X*_*i*_ might be reconstructed if *w* is known to the center. Consequently, our optimization strategy is able to preserve data privacy for all institutions. The whole framework of our federated Genotype-Expression-Image Integration model is summarized in **Algorithm 1**.

##### Algorithm 1. Federated Genotype-Expression-Image Data Integration Model.

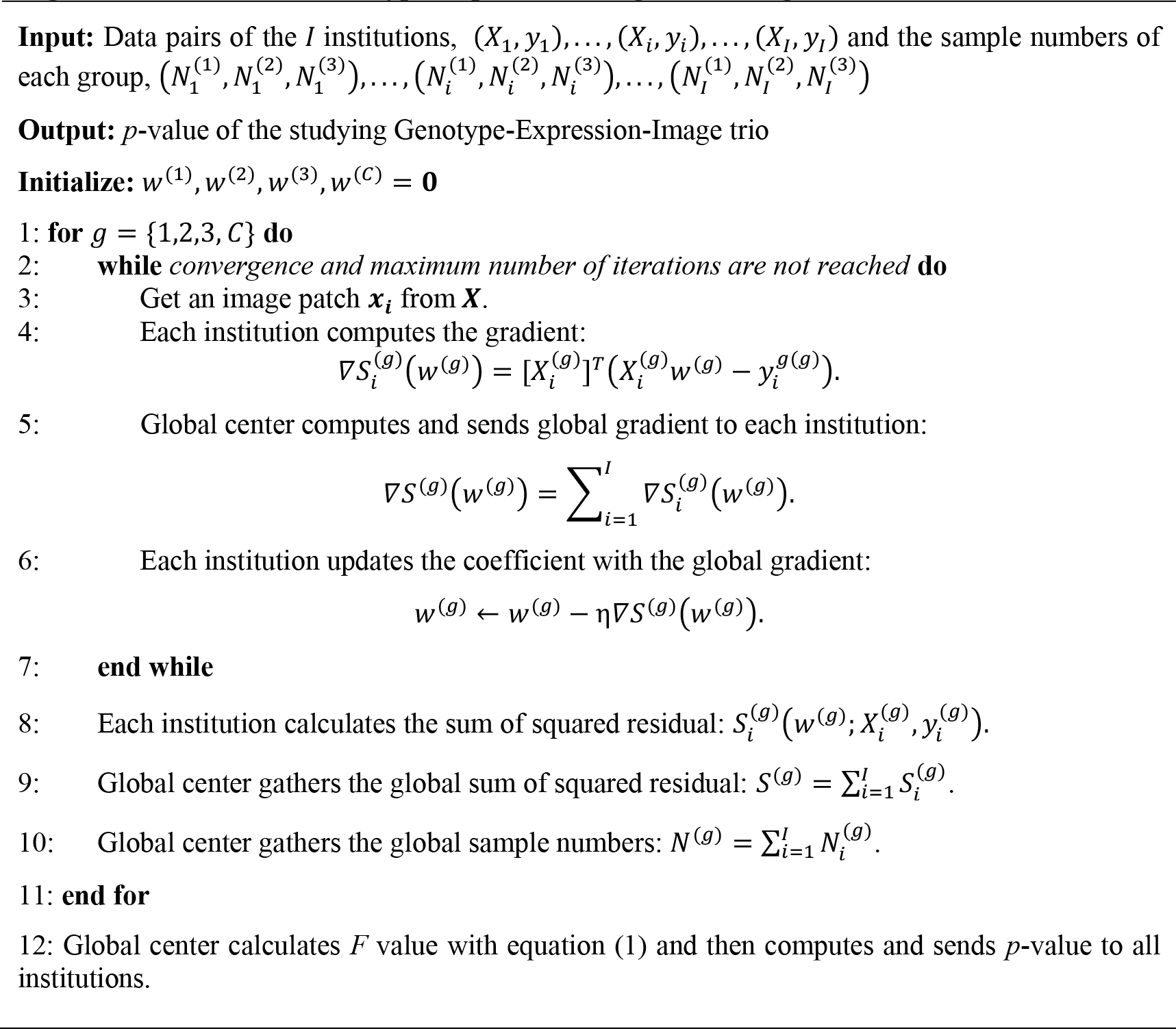

### 2.3 Performance Evaluation Protocol

We firstly use our model to identify AD-related gene expression. From the publicly available database, alzgene.org, we selected 619 known AD-related genes. When we fix the genotype and imaging biomarker, we can calculate a *p*-value for each of the 20,211 gene expressions. We rank the 20,211 *p*-values and identify which of the known AD-related genes are featured in the top *N* gene expressions. In Section 3.1, the top *N* gene expressions are the ones with a *p*-value < 0.05. In Section 3.4, *N* is a fixed number (100 and 200). We introduce hypergeometric analysis (Berkopec, 2007) to evaluate the model’s performance to detect the known AD-related genes. The probality mass function of hypergeometric analysis is defined as,

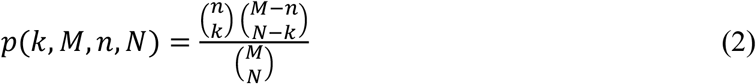

In our case, the number of population (*M*) is 20,211, the sample size (*n*) is 619, the number of samples drawn from the population (*n*) is the selected top *N* gene expressions, and the number of the observed successes (*k*) is the number of overlapping genes between 619 known AD-related genes and the top *N* gene expressions.

Secondly, with different genotypes, the pattern of hypergeometric enrichment will vary. The AD-related genotypes should, in general, have a more significant hypergeometric enrichment. From alzgene.org, we also obtain 1217 known AD-related SNPs. And we randomly select 1217 SNPs from the ADNI database as non-AD-related SNPs. After ranking the SNPs with the *p*-value based on hypergeometric analysis, we compute the number of AD-related SNPs found in the top *m* SNPs as true positive rate (TPR) and evaluate the performance of the models with TPR.

Finally, to prove the stability of our federated GEIDI, we compare the residuals of the federated linear regression model under different multi-site conditions. If the residuals are the same under different conditions, the *F* value and *p*-value will stay unchanged.

## 3 RESULTS

### 3.1 Discovering AD-related Gene Expressions

#### 3.1.1 APOE related Gene Expressions

*APOE* genotype is a well-known genetic biomarker for predicting subjects’ risk for AD. We stratify 697 subjects into three subgroups based on their *APOE* genotype status: non-carriers (e3/e3), heterozygotes (e3/e4), and homozygotes (e4/e4). Federated GEIDI is then adopted to discover genes correlated with hippocampus volume differentially across the three subgroups. We first run federated GEIDI with the volume of both sides of the hippocampus and the expression measures for 20,211 genes. Next, 1,625 gene expression measures are selected with *p* < 0.05. We evaluate the enrichment of these genes and the 619 AD-related genes annotated on alzgene.org and find 72 overlapping genes, yielding a hypergeometric enrichment *p* = 0.00036. Among the 72 overlapping genes, the top ten gene expressions were those measured for *CAST, CST3, GSTO1, LSS, MS4A4A, NPC1, PMVK, PPM1H, PPP2R2B, SORCS2*. Additionally, we performed the same experiments on the volume of the middle temporal gyrus (MidTemp); the results are shown in **Table 2**. 899 gene expressions are significant and 37 of them overlap with the 619 AD-related genes - with a hypergeometric enrichment *p* = 0.0053. The top ten gene expressions are those measured for *ABCA2, COL11A1, CST3, GNA11, HMOX1, HSPA1B, MAOA, MS4A4A, PRKAB2, SORCS2*.

**Table 2.** Hypergeometric statistics for *APOE*.

| Structures | Selected genes | Overlapping genes | $p$-value |
| --- | --- | --- | --- |
| Hippocampus | 1,625 | 72 | 0.00036 |
| MidTemp | 899 | 37 | 0.00528 |
| Linear Regression | 2,657 | 95 | 0.01236 |
| ANOVA | 3,234 | 107 | 0.02934 |

Matrix eQTL (Shabalin, 2012) is a state-of-the-art software to study the association between genotype and gene expression. We also leverage the linear model and ANOVA model in Matrix eQTL to evaluate the *APOE* genotype and the measured expression levels of the 20,211 genes. For the linear model, there are 2,657 significant gene expressions and 95 overlapping genes, leading to a hypergeometric enrichment *p* = 1.236*E* ‒ 02. For the ANOVA model, 3,234 gene expressions are selected, and 107 known genes are found, which leads to a *p*-value = 2.934*E* ‒ 02. The results show that our federated GEIDI can detect the most gene candidates that are significantly enriched for known AD genes. As the volume of hippocampus has the best performance in detecting AD-related genes, we use it as the imaging biomarker for all the remaining experiments.

#### 3.1.2 SNP related Gene Expressions

In this experiment, we stratify the subjects into three subgroups based on their SNP status. We choose *rs942439*, as this SNP is reported in alzgene.org, and use the volume of both sides of hippocampus as the imaging biomarker because of its superior performance in the first experiment. Federated GEIDI is used to detect any known AD gene whose expression is differentially associated with hippocampus volume in the subgroups stratified by the genotype at *rs942439* locus.

As shown in **Table 3**, 1,587 gene expressions were significant and 58 of them are reported in alzgene.org, leading to a hypergeometric enrichment p = 0.021. Of these 58 gene expression measures, the top ten genes are *ADRB1, ALOX5, ATXN1, CBS, FGF1, FLOT1, HSPA1A, RFTN1, SORL1, XRCC1*.

**Table 3.** Hypergeometric statistics for *rs942439*.

| Structures | Selected genes | Overlapping genes | *p*-value |
| --- | --- | --- | --- |
| **Hippocampus** | 1,587 | 58 | 0.021 |
| **Linear Regression** | 1,794 | 64 | 0.024 |
| **ANOVA** | 1,347 | 48 | 0.034 |

We also performed eQTL analysis on the SNP, *rs942439*. For a linear regression model, 1,794 gene expression values were selected and, of these, 64 genes are reported in alzgene.org, yielding a hypergeometric enrichment p = 0.024. For ANOVA model, For the linear regression model, 1,347 gene expression values were significant and, of these, 48 genes are reported in alzgene.org; in this case, the hypergeometric enrichment was p = 0.024.

In the experiment, one of the most significant gene expression measures is for *XRCC1*, for which the *p*-value is 4.332*E* ‒ 03. *XRCC1* is a gene coding for the X-ray repair cross-complementing protein; it was previously reported to be weakly associated with AD in a Turkish population (Doǧru-Abbasoǧlu et al., 2007).

As shown in **Figure 3**, we further adopt Pearson’s correlation to evaluate the relationship between the hippocampal volume (*x*-axis) (adjusted for ICV) and *XRCC1* gene expression (*y*-axis) of each subgroup. **Figure 3 (a)** illustrates the distribution for all the samples. **Figure 3 (b), (c) and (d)** show the distribution for the samples with “GG”, “GA” and “AA” genotype, respectively. Above each subfigure, R and p are the Pearson correlation coefficient and *p*-value, and N is the number of subjects. Even so, there is always some missing information in the genotype data. Hence, before we run federated GEIDI as well as the Pearson correlation statistics, we remove the subjects without the specific genotype. Because of this, the total number N in **Figure 3 (a)** is 579 instead of 697. We find samples with an “AA” genotype had hippocampal volume negatively correlated with expression levels of *XRCC1* (N=37, R=0.37, p=0.022). In contrast, the analysis in all samples (**Figure 3 (a))** or subjects with either “GG” or “GA” genotype (**Figure 3 (b), (c))** showed that the Pearson correlation coefficients were not significant in the overall, pooled sample. This result indicates that our method can establish associations among SNP, imaging, and gene expression data that include known AD risk factors.

**Figure 3.**
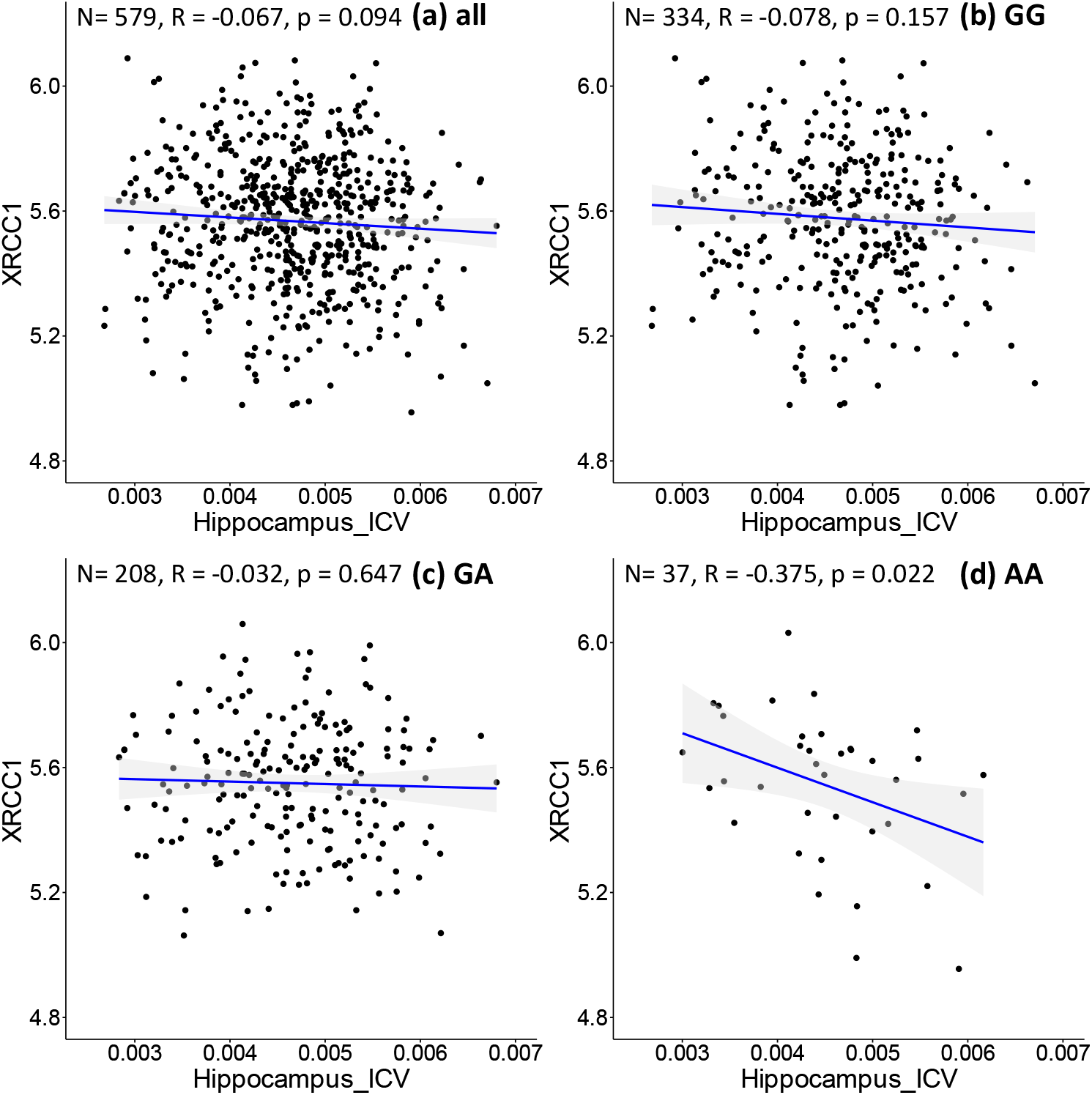
Correlation of image biomarkers and *XRCC1* gene expression in subpopulations stratified by the sample’s genotype at *rs942439*. **(a)** all samples **(b)** individuals with “GG” genotype; **(c)** those with “GA” genotype **(d)** those with “AA” genotype.

We further apply the above procedure to discover genes that have never been reported to be associated with AD. As shown in **Figure 4 (d)**, *SEC14L2* gene expression is negatively associated with hippocampal volume only in the subpopulation with “AA” genotype at rs942439 locus (N=37, R=-0.47, p=0.003). Interestingly, the opposite correlation is found in a subpopulation with “GA” genotype (**Figure 4 (c)**, N=208, R=0.15, P=0.03), and when applied to all pooled subjects, the total population does not show significant correlations (**Figure 4 (a)**, N=579, R=0.07, p=0.09). The *SEC14L2* gene encodes a protein that stimulates squalene monooxygenase, a downstream enzyme in the cholesterol biosynthesis pathway. This gene has never been reported to be associated with AD, but high cholesterol levels have been linked to early-onset AD (Wingo et al., 2019). This result indicates that our method can detect strong correlations in specific subpopulations that cannot be detected in the whole population. We also observe conflicting directions in different subpopulations, as shown by “GA” and “AA” subpopulations showing opposite correlations. This also highlights the importance of individualized medicine in patient management, as the same drug may have opposing effects in different groups of samples. Thus, federated GEIDI offers a new approach to discover novel genes related to AD as potential drug targets.

**Figure 4.**
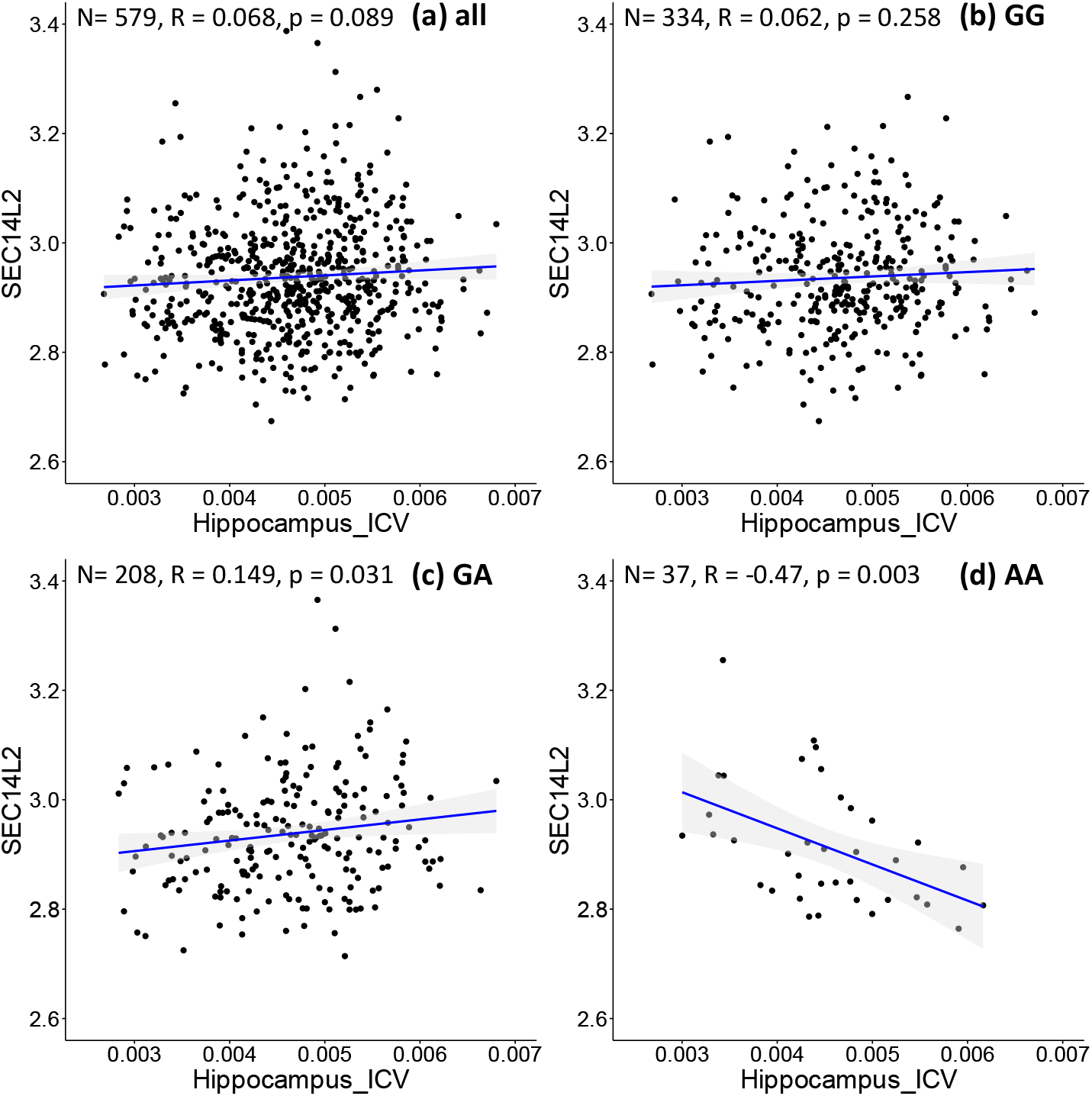
Correlation of image biomarkers and *SEC14L2* gene expression in subpopulation stratified by the sample’s genotype at *rs942439*. **(a)** all samples **(b)** those with “GG” genotype; **(c)** those with “GA” genotype; **(d)** those with “AA” genotype.

### 3.2 Discovering AD-related SNPs

In the experiments of **Section 3.1**, we used hypergeometric statistics to evaluate the ability of our proposed model to discover AD-related gene expressions that are differentially associated with imaging measures in populations stratified by *APOE* haplotype. In this experiment, we also use hypergeometric statistics to assess the discovery rate of known AD-related genes, in the set of genes whose expression shows different correlations with imaging markers, in samples stratified according to different genotypes. Sets that are enriched in AD-related SNPs will have a more significant *p*-value in the hypergeometric test that assesses enrichment. Since the hippocampal volume measure showed superior performance for this task, among all the imaging biomarkers in Sec. 3.1, we adopt it as the brain imaging measure in this experiment. To illustrate the effectiveness of our GEIDI model, we perform the same experiment with the linear model in Matrix eQTL, which can evaluate the associations between SNPs and gene expression.

When we analyze each SNP with our federated GEIDI and Matrix eQTL, we will obtain a *p*-value for each of the 20,211 expressed genes. Instead of selecting the significant gene expressions with a *p*-value < 0.05, we respectively rank the *p*-value of all the gene expressions calculated by the two methods and select the top *N* (100 and 200) gene expressions to apply the hypergeometric analysis. With the *p*-value from this hypergeometric analysis (which assesses enrichment for known AD-associated genes), we may rank the SNPs and obtain the most AD-related ones. Then, we try to prove that our GEIDI is able to detect more AD-related SNPs. From AlzGene.org, we also created a list of 1,217 AD-related SNPs, and we randomly selected another 1,217 SNPs as the non-AD-related ones. After ranking the SNPs with the *p*-value computed by the two methods, we calculate the true positive rate (TPR) for the top *m* SNPs, which measures the percentage of AD-related SNPs in the selected top *m* SNPs. For example, the last number in **Table 4** is 0.57, which means 57% of the top 500 SNPs are AD-related ones. As the results in **Table 4**, our federated GEIDI can always achieve superior performance than Matrix eQTL.

**Table 4.** True Positive Rate of AD-related SNPs in the top *m* SNPs.

| Matrix eQTL: Linear Regression |  |  |  |  |  |
| --- | --- | --- | --- | --- | --- |
| SNP ($m$) \ EXP ($N$) | 10 | 50 | 100 | 200 | 500 |
| 100 | 0.50 | 0.58 | 0.54 | 0.54 | 0.53 |
| 200 | **0.50** | 0.60 | 0.57 | **0.58** | 0.54 |
| Matrix eQTL: ANOVA |  |  |  |  |  |
| SNP ($m$) \ EXP ($N$) | 10 | 50 | 100 | 200 | 500 |
| 100 | 0.60 | 0.56 | 0.52 | 0.50 | 0.53 |
| 200 | **0.50** | 0.60 | 0.57 | 0.55 | 0.56 |
| Federated GEIDI |  |  |  |  |  |
| SNP ($m$) \ EXP ($N$) | 10 | 50 | 100 | 200 | 500 |
| 100 | **0.60** | **0.60** | **0.61** | **0.61** | **0.60** |
| 200 | **0.50** | **0.62** | **0.61** | **0.58** | **0.57** |

The SNPs are ranked with the $p$ -value from hypergeometric analysis with the top $N$ gene expressions as the number of samples drawn from the population.

In **Figure 5**, we visualized the *p*-values of these 2,434 SNPs from hypergeometric analysis in the Manhattan plots.. The top figure is the Manhattan plot for the result with the top 100 gene expressions and the bottom one is for the result of the top 200 gene expressions. The SNPs, *rs4889013* and *rs11940059*, are the top-ranked ones for both results. When we select 100 or 200 as the number of samples drawn from the population, three parameters in **Equation (2)** are fixed and only the number of observed successes, *k*, varies for different SNPs. Therefore, the *p*-value from different SNPs might be the same if their numbers of observed successes are the same. This explains why some SNPs locate at the same horizontal position.

**Figure 5.**
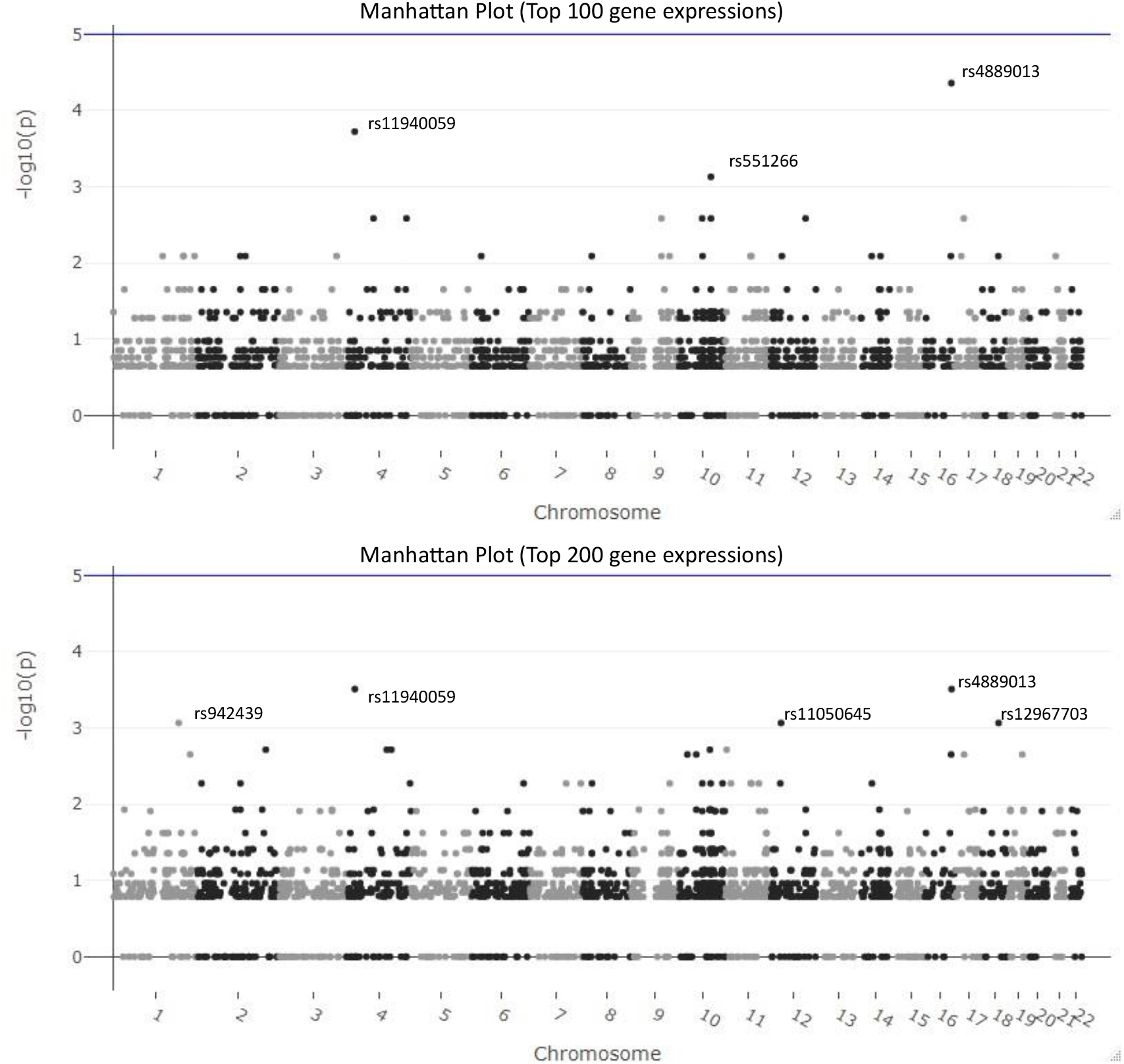
Manhattan plots for the results of federated GEIDI. The top figure is the Manhattan plot for the results from hypergeometic analysis with the top 100 gene expressions as the number of samples drawn from the population and the bottom one is for the results with the top 200 gene expressions. The SNPs, *rs4889013* and *rs11940059*, are the top-ranked ones for both results.

### 3.3 Federated Learning Stability Analysis

In this experiment, we aim to demonstrate that the performance of our federated GEIDI model is not greatly affected by different data distribution models across institutions. In practice, it would be convenient and efficient to run association tests on data that might be distributed across multiple servers, without having to transfer it all to a centralized location. We synthesized 1,000 samples and randomly assigned them to different independent hypothetical institutions, including one institution, three institutions, five institutions and seven institutions. We compared the residuals from each linear regression model for each condition and found the residuals remained unchanged, as shown in **Table 5**. The first column is the ground truth residual and the rest are the residuals for our federated linear model under different data distribution conditions. The residuals are the same, which means that the results of our Federated GEIDI will remain stable under different multi-site conditions. Therefore, these results demonstrate the correctness and stability of our federated GEIDI model.

**Table 5.**
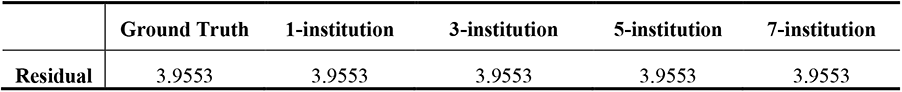
Stability analysis of federated GEIDI across different institutional settings.

## 4 DISCUSSION

In this work, we propose a novel federated Genotype-Expression-Imaging Data Integration (GEIDI) model to identify the genetic and transcriptomic influences on brain sMRI measures. We performed various experiments with our model on the publicly available ADNI dataset, and we have two main findings. First, our federated GEIDI is an effective multimodal approach that provides novel insights into the relationship among image biomarkers, genotypes, and gene expression, and may be useful to discover novel genes as potential AD drug targets. It has better performance in detecting AD-related gene expressions and SNPs than the linear regression model and ANOVA model in the state-of-the-art Matrix eQTL approach. In addition, our model may not only detect known AD-associated genes as potential drug targets, such as *XRCC1*, but may also help in discovering novel genes as potential drug targets, such as *SEC14L2*. Second, compared to Matrix eQTL, our federated GEIDI provides a way to investigate extremely large datasets from different institutions without violating data privacy. The statistical power of the model will also be increased with the larger sample size. Our work may lay down a solid foundation for future multi-site large-scale imaging genetics research.

### 4.1 Comparison Analysis of Federated GEIDI and Matrix eQTL

Expression quantitative trait loci (eQTL) analysis (Nica and Dermitzakis, 2013; Rockman and Kruglyak, 2006) is designed to identify the significant associations between SNPs and gene expression, which can help understand the biochemical processes occurring in living systems, discover the genetic factors that influence the onset and progression of certain diseases, and determine the pathways affected by them. There are many eQTL analysis methods, including linear regression, ANOVA models, Bayesian regression (Servin and Stephens, 2007), and so on. Matrix eQTL (Shabalin, 2012) is the state-of-the-art software for computationally efficient eQTL analysis, and it supports additive linear and ANOVA models. It has been widely used in the study of human genetic traits and diseases. However, it has two main limitations. First, although Matrix eQTL is very computationally efficient, it cannot work on data that is distributed across different institutions. Nowadays, unprecedentedly large volumes of biomedical and genetic data have been collected by different hospitals and research institutions, and this aggregate of available data may significantly advance the study of factors influencing disease. However, data restrictions, legal complexities, and patient privacy have all been major obstacles for researchers to obtain or share these data. Therefore, federated machine learning and distributed statistical models are becoming advantageous for current research on medical data (Ng et al., 2021; Sheller et al., 2020). Second, the models in Matrix eQTL cannot jointly consider the information from images. Changes in brain structures can play a vital role in the study and diagnosis of Alzheimer’s disease, and many researchers have attempted to detect associations between genetic factors and imaging features (Dong et al., 2019; Shen et al., 2020; Shen and Thompson, 2020). Therefore, introducing imaging information may greatly assist the detection of genetic factors that influence disease as an intermediate phenotype that might reflect relevant disease processes.

Our proposed federated Genotype-Expression-Imaging Data Integration model can effectively overcome these two obstacles. In the Methods section, we detailed how our model maintains each institutional data private. Additionally, our federated GEIDI model integrates GWAS data, gene expression, and imaging data. The experimental results demonstrate that our federated GEIDI model has better performance in detecting AD-related genes and SNPs. In detecting AD-related gene expression, our model achieves the strongest hypergeometric enrichment with the volume of the hippocampus. In our tests detecting AD-related SNPs, our federated GEIDI model generally obtained a higher TPR than the linear regression model and ANOVA model. Besides, compared with existing methods, our proposed model offers novel insights into the relationship among image biomarkers, genotypes, and gene expression by considering both imaging and gene expression features – which can vary over time - and understanding how they are affected by an individual’s SNPs.

### 4.2 Drug Target for Precision Medicine of AD

Increasingly, a major challenge in healthcare is that many drugs are adequate for only small subgroups of patients (Schork, 2015). Some patients may not only suffer from adverse side effects but also waste money on ineffective drugs. Precision medicine has the potential to tailor therapy based on the best expected response and highest safety margin to ensure better patient care. By enabling each patient to receive earlier diagnoses, risk assessments, and optimal treatments, personalized medicine holds promise for improving health care while also potentially lowering costs (Vogenberg et al., 2010). In this work, our multi-omics approach offers potential in genome-guided drug discovery. Compared to state-of-the-art methods, our model performs better in detecting AD-related genes and SNPs. Moreover, our model cannot only detect known genes for target drugs, like *XRCC1*, but also can discover novel potential gene expressions, like *SEC14L2*. Meanwhile, our federated framework may integrate data from multiple sourses without violating the data praicay and the larger sampe size will help discover and understand more AD-related genetic information. Consequentially, we believe our federated GEIDI model will play an important role in the study of precision medicine in the future.

### 4.3 Limitations and Future Work

Despite the promising results of our federated GEIDI model, there are three caveats. Firstly, we only evaluated our model on data from 697 subjects from the publicly available ADNI dataset. In the future, we will add other datasets to make results more robust and reliable. For example, the Arizona APOE cohort (AZ APOE cohort) recruited 450 actively followed participants matched by age, sex, and education – including homozygous *APOE-e4* carriers and non-*e4* carriers since 1994 (Caselli et al., 2009). The UK Biobank project (Cox, 2018) collects both large-scale genetic-genomic and phenotypic data as well as health-related information from around 500,000 volunteer participants in the UK. Assessments include biological measures, blood- and urine-based biomarkers, body, and brain imaging scans, and lifestyle parameters (Bycroft et al., 2018; Elliott et al., 2018). Second, the volumes of specific subcortical structures may not be ideal imaging measurements for the multiple biological processes involved in Alzheimer’s disease. Surface-based morphometry analyses have achieved excellent performance for early AD detection (Wang et al., 2010; Wu et al., 2018; Zhang et al., 2017). In recent work (Wang et al., 2021; Wu et al., 2021), the authors created tools to generate a univariate morphometry index (UMI) for surface morphometry features on regions of interest (ROIs) that are related to beta-amyloid deposition. This induced UMI may reflect intrinsic morphological changes induced by processes of amyloid accumulation in AD. and have greater signal-to-noise ratio and strong generalizability to new subjects. If we were to use such brain pathology induced UMI measures instead of volumes, our federated GEIDI model may detect additional AD-related genes whose expression is influenced by SNPs. Finally, in ongoing work on blood-based biomarkers (Bateman et al., 2019; Janelidze et al., 2020), plasma levels of amyloid-beta (plasma Aβ) may provide an alternative but highly accurate estimate of brain amyloid positivity. In (Janelidze et al., 2020), plasma P-Tau181 accurately discriminated AD dementia from non-AD neurodegenerative diseases with an excellent AUC (0.94). Similarly, such plasma measures might be used in conjunction with our federated GEIDI model to better understand effects of AD-related genotypes. We plan to analyze such datasets to further evaluate our model in the future.

## 5 CONCLUSION

We propose a novel federated Genotype-Expression-Image Data Integration model. Compared to similar studies, this work achieves state-of-the-art performance in discovering downstream effects AD-related genes and SNPs. Besides, the model provides novel insights into the relationship among image biomarkers, genotypes, and gene expression and could discover novel drug targets for percision medicine. In the future, we will further validate our model with more datasets and more advanced imaging biomarkers. Specifically, we will introduce blood-based biomarkers into our model when such data are available.

## ACKNOWLEDGMENTS

Algorithm development and image analysis for this study were partially supported by the National Institute on Aging (RF1AG051710, R21AG065942, U01AG068057, R01AG031581, and P30AG19610), the National Institute of Biomedical Imaging and Bioengineering (R01EB025032), Mayo-ASU seed funding, and National Library of Medicine (R01LM013438).

Data collection and sharing for this project was funded by the Alzheimer’s Disease Neuroimaging Initiative (ADNI) (National Institutes of Health Grant U01 AG024904) and DoD ADNI (Department of Defense award number W81XWH-12-2-0012). ADNI is funded by the National Institute on Aging, the National Institute of Biomedical Imaging and Bioengineering, and through generous contributions from the following: Alzheimer’s Association; Alzheimer’s Drug Discovery Foundation; BioClinica, Inc.; Biogen Idec Inc.; Bristol-Myers Squibb Company; Eisai Inc.; Elan Pharmaceuticals, Inc.; Eli Lilly and Company; F. Hoffmann-La Roche Ltd and its affiliated company Genentech, Inc.; GE Healthcare; Innogenetics, N.V.; IXICO Ltd.; Janssen Alzheimer Immunotherapy Research & Development, LLC.; Johnson & Johnson Pharmaceutical Research & Development LLC.; Medpace, Inc.; Merck & Co., Inc.; Meso Scale Diagnostics, LLC.; NeuroRx Research; Novartis Pharmaceuticals Corporation; Pfizer Inc.; Piramal Imaging; Servier; Synarc Inc.; and Takeda Pharmaceutical Company. The Canadian Institutes of Health Research is providing funds to support ADNI clinical sites in Canada. Private sector contributions are facilitated by the Foundation for the National Institutes of Health (www.fnih.org). The grantee organization is the Northern California Institute for Research and Education, and the study is coordinated by the Alzheimer’s Disease Cooperative Study at the University of California, San Diego. ADNI data are disseminated by the Laboratory for Neuro Imaging at the University of Southern California.

